# Short-term methionine starvation induces de novo diurnal oscillations of hepatic m6A RNA methylation

**DOI:** 10.64898/2026.07.03.736420

**Authors:** Yang Liu, Kaliopi Chrysovergis, Katina L. Johnson, Jason G. Williams, Fred B. Lih, Leesa J. Deterding, Sara A. Grimm, Paul A. Wade

## Abstract

Dietary methionine restriction has been shown to improve metabolic health and treat multiple diseases. Methionine metabolism regulates transmethylation reactions, including N6-methyladenosine (m6A) RNA methylation, by modulating the availability of S-adenosyl methionine (SAM). Both m6A RNA methylation and methionine metabolism are involved in the regulation of the circadian clock. However, it remains unclear whether dietary methionine influences circadian rhythms through the regulation of m6A RNA modification. In this study, we investigated the effects of short-term methionine deprivation on the diurnal oscillations of m6A RNA methylation in the mouse liver. We found that a methionine-deficient (MD) diet reprogrammed the cyclic expression patterns of m6A writers, erasers, and readers. Methylated RNA immunoprecipitation sequencing (MeRIP-seq) revealed that the MD diet induced de novo diurnal m6A oscillations in genes associated with RNA processing, protein translation, protein ubiquitination, and mTORC1 signaling pathways. RNA-seq and quantitative proteomics analyses demonstrated that MD-induced changes in m6A RNA levels were linked to alterations in mRNA and protein abundance. We observed that dynamic m6A RNA methylation of the transcripts encoding two key enzymes, MAT2A and CBS, helps maintain methionine homeostasis in response to methionine starvation. These findings identify m6A RNA methylation as a key mechanism linking methionine metabolism to circadian regulation.

## INTRODUCTION

Methionine metabolism influences various cellular processes, including epigenetic regulation, maintenance of redox balance, protein translation, and lipid homeostasis ^1,2^. Dietary methionine restriction (MR) has been shown to improve metabolic health, extend lifespan, suppress tumor growth, and enhance cancer treatment ^3–6^. Interestingly, accumulating evidence demonstrates that methionine metabolism plays a role in the regulation of circadian rhythms ^7–9^. Our recent study demonstrated that short-term methionine starvation leads to dramatic reprogramming of hepatic circadian rhythms ^10^. These studies provide evidence supporting the notion that nutritional challenges may regulate various biological processes in a circadian-dependent manner.

m6A RNA modification is the most abundant internal modification in eukaryotic RNAs, playing significant roles in regulating RNA splicing, translation, and stability ^11^. This modification influences circadian rhythms by regulating RNA processing of several core clock genes, including *Per2* and *Arntl* ^12^. It also modulates the translation of casein kinase 1 delta (CK1δ), a key kinase involved in circadian clock control ^13^. A recent study shows that m6A methylation of *Nr1d1* leads to mRNA degradation, disrupting the circadian clock in hepatic stellate cells (HSCs) ^14^. Notably, liver-specific deletion of the m6A writer METTL3 impairs the nuclear export of mRNAs from core clock genes, such as *Arntl* and *Clock* ^15^. These findings underscore the crucial role of m6A RNA modification in the circadian regulation network.

SAM, synthesized by methionine adenosyltransferases (MATs) from methionine and adenosine triphosphate (ATP), serves as the primary methyl donor during transmethylation of substrates, including DNA, RNA, and proteins. SAM abundance can be sensed by its protein sensors, such as SAMTOR and PRMT1, which regulate the downstream mTORC1 pathway ^16,17^. m6A methyltransferase 16 (METTL16), a key regulator of SAM homeostasis, has also been proposed to function as an intracellular SAM sensor ^18,19^. When SAM levels are adequate, METTL16 binds to and methylates MAT2A in the 3’ untranslated region (UTR). Upon SAM depletion due to methionine starvation, reduced m6A RNA methylation leads to increased mRNA stability and pre-mRNA splicing, which in turn induces MAT2A expression to compensate for decreased SAM levels ^18^. In addition to the m6A levels of *Mat2a*, a recent study revealed that MR affects immunotherapy responses by modulating m6A RNA modification in tumor cells ^20^. These studies provide evidence supporting the close link between methionine metabolism and m6A RNA modification. Although both methionine metabolism and m6A RNA modification have been implicated in circadian regulation, whether methionine starvation regulates the circadian clock through m6A RNA methylation remains unknown.

In this study, we performed MeRIP-seq to explore the effects of short-term methionine starvation on m6A RNA methylation and found that the MD diet induced the formation of de novo m6A methylation in the liver. Furthermore, RNA-seq and quantitative proteomics confirmed that MD diet-induced m6A alterations were associated with changes in mRNA and protein abundance. These findings demonstrate that m6A RNA modification plays an important role in the regulation of methionine metabolism and the circadian clock, providing insights into the molecular mechanisms underlying the nutritional regulation of the circadian rhythms.

## Results

### Altered circadian oscillations of genes involved in m6A RNA methylation by the MD diet

To investigate the effects of dietary methionine on circadian rhythms, six-month-old male C57BL/6J mice were placed on either a control diet containing 0.82% methionine or an MD diet with no methionine for three weeks (Figure 1A). Serum was collected from the mice at four time points, and the relative abundance of methionine and SAM was measured. We observed that methionine starvation led to a dramatic reduction in serum methionine levels. Additionally, we found that methionine and SAM levels at zeitgeber time (ZT) 21 were significantly lower compared to those at ZT9, only in mice on the MD diet. In contrast, no oscillations in methionine and SAM levels were detected in mice on the control diet (Figure 1B), suggesting that short-term methionine deprivation alters the circadian rhythms of methionine metabolism.

**Figure 1.**
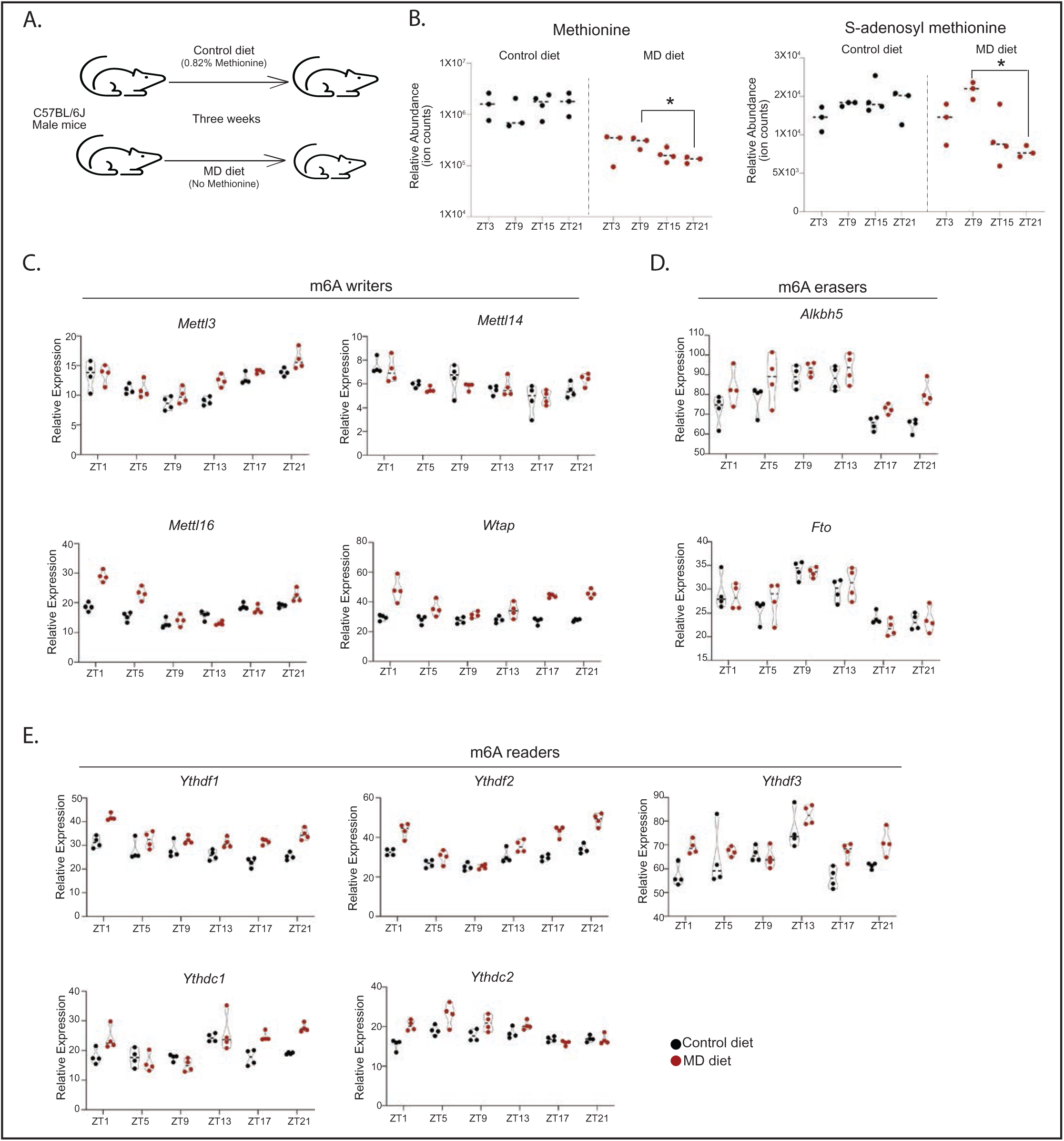
Short-term methionine starvation alters the circadian oscillations of genes involved in m6A RNA methylation. A) Diagram showing that six-month-old male C57BL/6J mice were placed on either a control diet with 0.82% methionine or an MD diet for three weeks. B) The relative abundance of methionine and S-adenosyl-methionine in serum collected from mice on the indicated diet at four time points. (*p < 0.05, one-way ANOVA, Tukey’s multiple comparisons test). C-E) TMM-normalized gene expression values of m6A writers (C), erasers (D), and readers (E) in the liver of mice fed the indicated diet for three weeks.

To assess the impact of the MD diet on the oscillation of m6A RNA modification, we examined the mRNA expression of genes involved in m6A methylation by analyzing RNA-seq data from mouse liver. The results revealed that the expression of the m6A “writer” genes *Mettl3* and *Mettl14* remained largely unchanged by the MD diet, except that Mettl3 expression increased at ZT13. However, the levels of *Mettl16* were increased by the MD diet at ZT1 and ZT5, and the expression of *Wtap* was elevated at ZT1, ZT17 and ZT21 (Figure 1C). Regarding the m6A “eraser,” we observed that the expression of *Alkbh5* was increased at ZT21 in mice on the MD diet, while the expression of *Fto* remained unchanged (Figure 1D). We also analyzed the expression levels of five m6A “reader” genes. The MD diet led to an increase in the expression of *Ythdf1*, *Ythdf2*, and *Ythdf3* at ZT1, ZT17, and ZT21. Additionally, *Ythdc1* expression was elevated at ZT17 and ZT21, and *Ythdc2* expression was higher at ZT1 in mice on the MD diet compared to those on the control diet (Figure 1E). These findings demonstrate that the MD diet reshapes the oscillating expression of m6A RNA methylation-related genes, suggesting that methionine regulates the circadian rhythms of m6A RNA methylations.

### Hepatic diurnal oscillations of m6A RNA methylation in response to methionine starvation

To explore the effects of methionine starvation on m6A RNA modification, we assessed the m6A methylation of mRNA by performing MeRIP-seq on liver tissues collected from mice on a control or MD diet at ZT9 and ZT21. We identified 9,753 m6A methylation peaks and generated metagene plots with these peaks (Table S1). Consistent with previous studies ^21,22^, m6A methylation was highly enriched in the vicinity of the stop codon in the liver and predominantly occurred within the RRACH consensus motif (Figure 2A and 2B).

**Figure 2.**
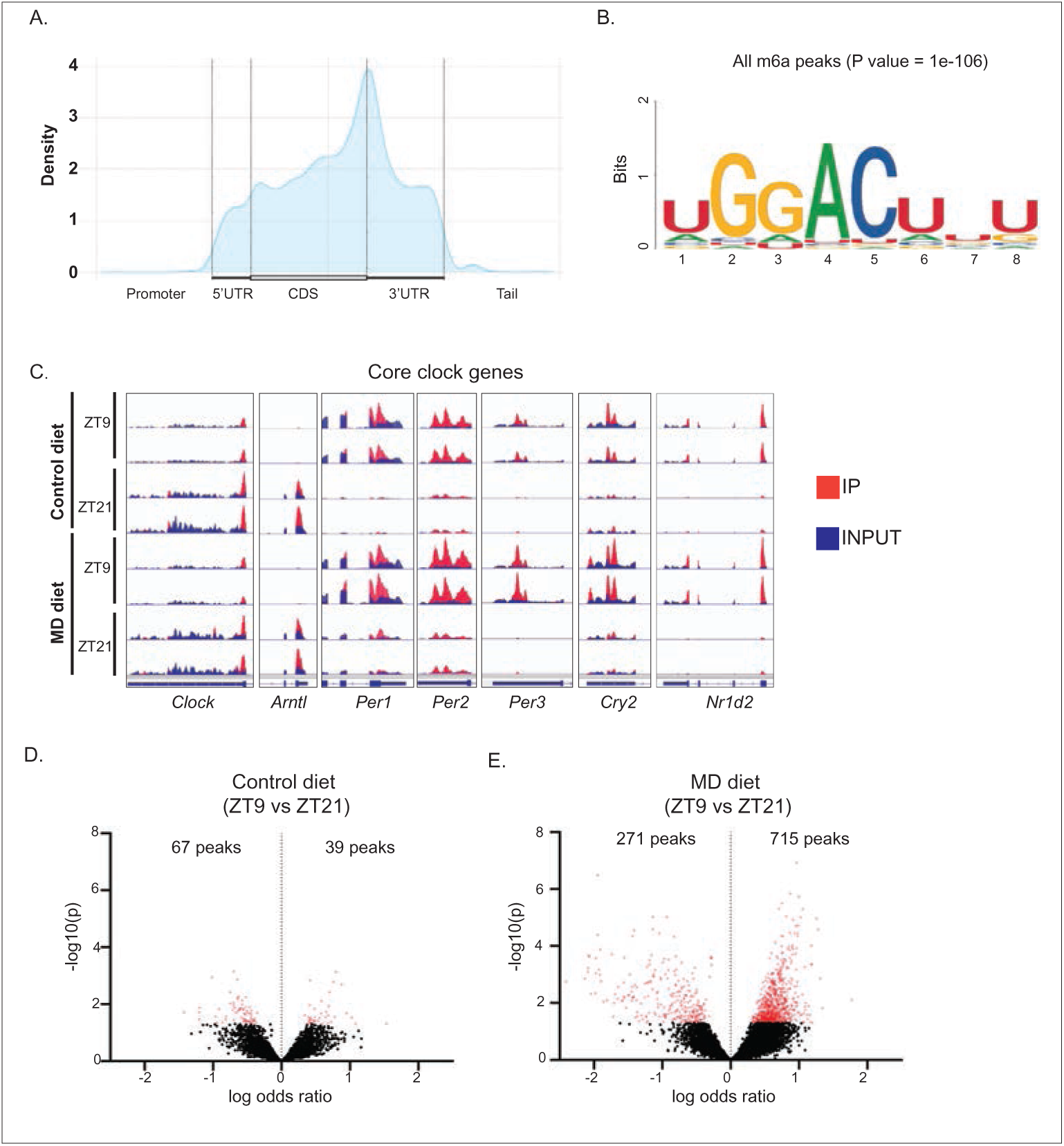
De novo diurnal oscillations of m6A RNA modifications induced by methionine starvation. A) The metagene plot depicts the density distribution of m6A in the liver as detected by MeRIP-seq. Position within the transcript is shown on the x-axis, and density is shown on the y-axis. B) Motif enrichment in RIP-seq peaks was determined by HOMER using all m6A peaks. C) Integrative Genomics Viewer (IGV) browser tracks showing m6A RNA methylation in the mRNA of core clock genes. D-E) Volcano plots depict transcripts exhibiting differential m6A RNA methylation, as measured by RIP-seq, comparing ZT9 to ZT21 in the liver of mice fed the control diet (D) and MD diet (E).

We then investigated the effects of the MD diet on the rhythmicity of m6A methylation in the liver. Although m6A methylation was observed in several core clock genes, including *Clock*, *Arntl*, *Per1*, *Per2*, *Per3*, *Cry2*, and *Nr1d2*, we found no significant changes in m6A methylation in these genes between ZT9 and ZT21 under both Control and MD conditions (Figure 2C). Notably, only ∼100 m6A peaks exhibited rhythmic methylation in mice on the control diet (Figure 2D and Table S2).

In contrast, under the MD condition, approximately 1,000 m6A peaks showed significant differences in methylation levels between ZT9 and ZT21 (Figure 2E and Table S3). Among these, 715 m6A peaks exhibited lower methylation levels at ZT21 compared to ZT9. These findings suggest that methionine starvation reshapes the oscillations of m6A RNA modification.

We conducted gene ontology analysis for 645 genes with reduced m6A methylation at ZT21 compared to ZT9 in mice (Table S4). We found that these genes were enriched in processes such as protein ubiquitination, translation, protein transport, mRNA splicing and processing, and the mammalian target of rapamycin complex 1 (mTORC1) signaling pathway (Figure 3A). The m6A peaks in the 3’ UTR of genes, including *Rptor*, *Mfsd8*, and *Mlst8*, which are involved in mTORC1 pathways, were reduced at ZT21 compared to ZT9, specifically in mice on the MD diet. This suggests that methionine metabolism may control the oscillations of the mTORC1 pathway via modulation of m6A RNA modification (Figure 3B). We also found that the MD diet induced m6A oscillations in genes related to protein ubiquitination, including ubiquitin-conjugating enzymes (*Ube2d2a* and *Ube2g1*) and ubiquitin ligases (*Rnf7*, *Rbx1*, *Nedd4*, and *Trim27*) (Figure 3C). Interestingly, recent studies have demonstrated that the mRNA stability of several ubiquitin ligases, including *Nedd4*, *Trim27*, and *Rnf7*, is regulated by m6A RNA methylation ^23–25^. Genes involved in protein translation, such as *Mrpl4*, *Mrps17*, *Mrpl10*, *Mrps23*, *Etf1*, and *Rsl24d1*, also exhibited reduced m6A levels at ZT21 compared to ZT9, specifically in mice fed the MD diet (Figure 3D). Taken together, our findings suggest that de novo m6A oscillations induced by the MD diet may regulate protein and RNA homeostasis via modulating RNA splicing, protein degradation and translation.

**Figure 3.**
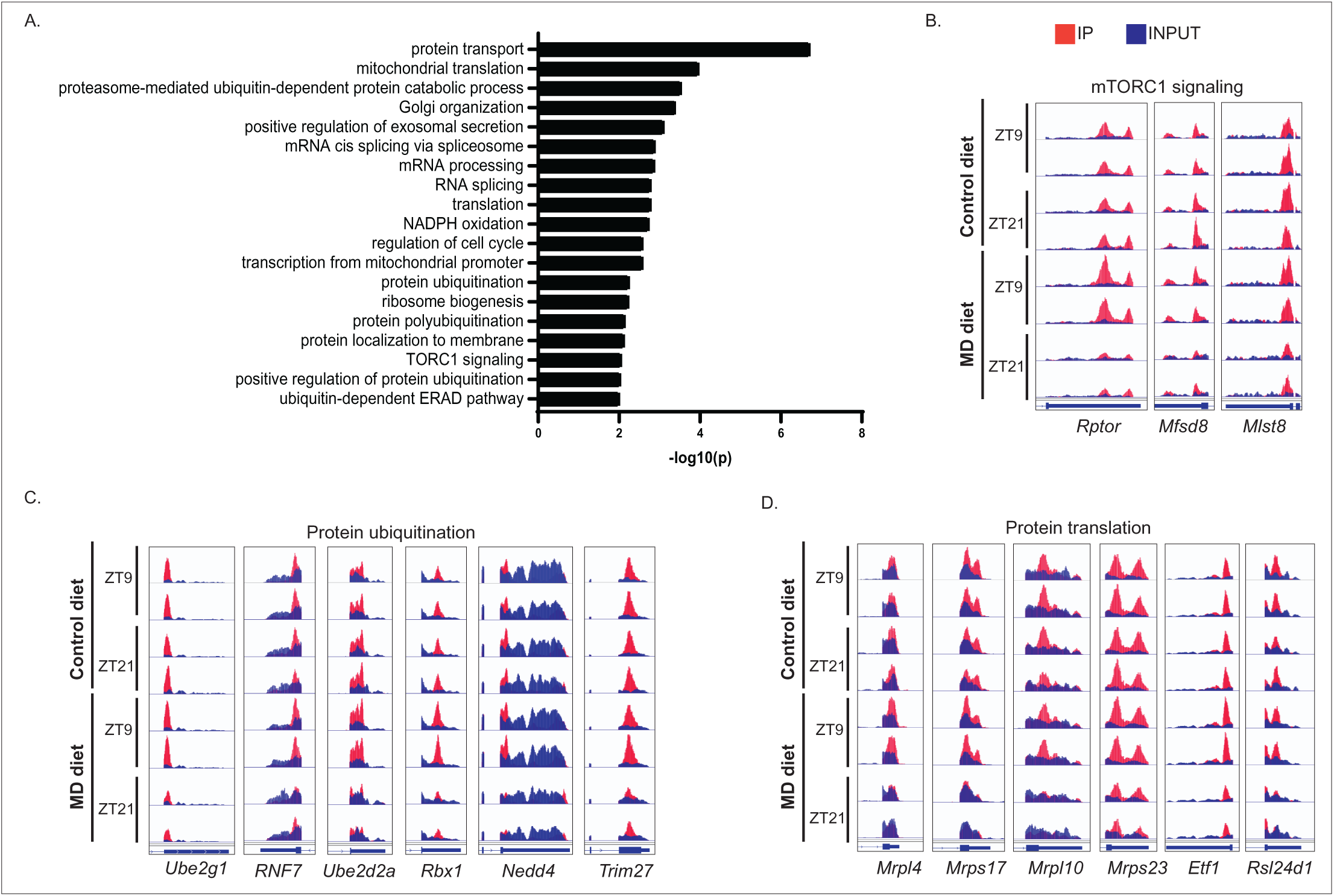
Pathway analysis of genes with reduced m6A RNA methylation at ZT21 in mice fed the MD diet. A) Gene ontology analysis was conducted on transcripts with reduced m6A RNA methylation at ZT21 compared with ZT9 in animals on the MD diet. The bar graph indicates enriched GO terms, with p-values shown on the axis. B-D) IGV browser tracks showing m6A RNA methylation of genes in the mTORC1 signaling (B), protein ubiquitination (C), and protein translation (D) pathways.

### Changes in RNA and protein levels associated with MD diet-induced alterations of m6A mRNA modifications

Given that m6A RNA methylation regulates mRNA stability and protein translation pathways in response to methionine starvation, we aimed to determine whether changes in m6A methylation induced by the MD diet were associated with alterations in mRNA and protein levels. Compared to the m6A mRNA methylation in control diet-fed mice, we identified 950 m6A peaks with significantly differential methylation in MD-fed mice at ZT21 (Figure 4A and Table S5). Among these, methylation levels at 726 peaks, including in the mRNAs of *Mat2a*, *Cbs*, and *Ddx17*, were reduced by the MD diet (Figure 4A). Furthermore, these m6A sites with reduced methylation in MD-fed mice were enriched in regions with the RRACH motif (Figure 4B).

**Figure 4.**
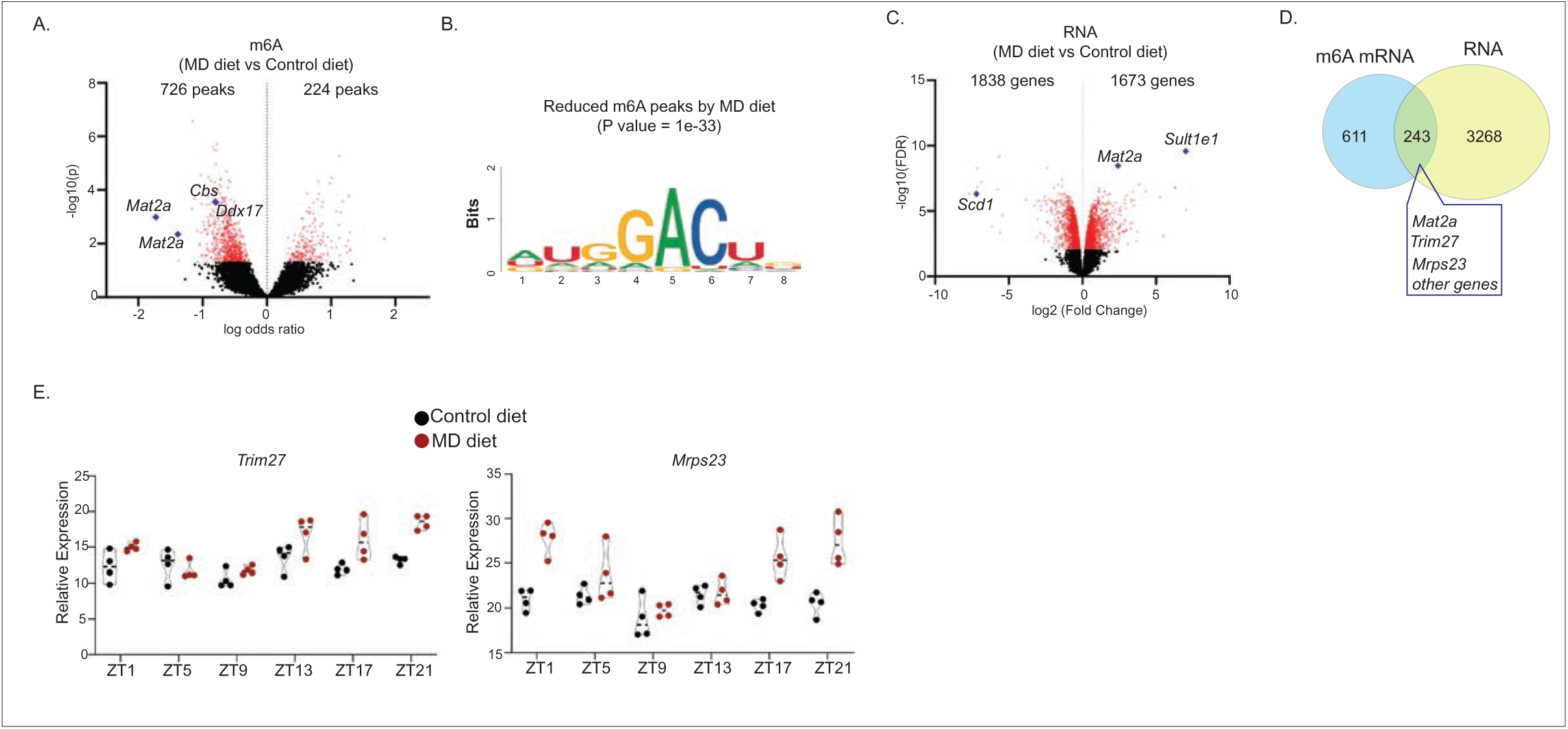
Alterations in m6A RNA modifications associated with changes in RNA upon MD diet. A) Volcano plot depicting transcripts exhibiting differential m6A RNA methylation as measured by RIP-seq in the liver induced by the MD diet at ZT21. B) Motif enrichment was determined as described above in the 726 peaks with reduced m6A methylation at ZT21 compared with ZT9 in MD-fed mice. C) Volcano plot depicting differentially expressed genes (DEGs) in the liver induced by the MD diet at ZT21. D) Venn diagram depicting the overlap between transcripts with differential m6A RNA methylation and DEGs in the liver at ZT21. E) TMM-normalized gene expression values of *Trim27* and *Mrps23*.

We analyzed RNA-seq data from livers collected at ZT21. Compared to the control diet, we identified 3,511 differentially expressed genes (DEGs, with FDR < 0.01) in MD diet-fed mice (Figure 4C and Table S6). Of these, 1,673 genes, including *Mat2a* and *Sult1e1*, showed higher expression levels in MD diet-fed mice, while the expression of 1,838 genes, including *Scd1*, was downregulated by the MD diet (Figure 4C). Notably, we found that m6A methylation levels of 243 DEGs, including *Mat2a*, *Trim27*, and *Mrps23*, were altered by the MD diet (Figure 4D and Table S7). In contrast to the reduced m6A methylation levels of *Trim27* and *Mrps23* in MD-fed mice at ZT21, the expression of these two genes was increased by the MD diet at ZT21(Figures 4E), suggesting that m6A may influence their mRNA stability.

To investigate the effects of the MD diet on protein expression, we performed quantitative proteomic analysis using liver tissues collected at ZT21. We identified 635 proteins that were significantly altered (adjusted p < 0.05) between the control and MD diet groups (Figure 5A and Table S8), including 280 proteins with reduced abundance and 355 proteins with increased abundance in the livers of MD diet-fed mice. Overall, protein abundance changes were positively correlated with mRNA expression changes (Figure 5A). For example, both mRNA and protein levels of SCD1 were markedly decreased, whereas SULT1E1 levels were increased in response to the MD diet. However, several proteins, including CBS and DDX17, exhibited altered protein abundance despite minimal changes in their corresponding mRNA levels, suggesting potential post-transcriptional regulation. Notably, we identified 62 proteins, including MAT2A, CBS, and DDX17, that displayed both differential m6A RNA methylation of their transcripts and altered protein abundance in response to the MD diet (Figure 5B and Table S9). DDX17, a member of the large family of DEAD-box RNA helicases, plays a critical role in various RNA functions ^26^. The MD diet led to reduced m6A methylation and increased protein levels of DDX17, without affecting its mRNA levels (Figures 5A-D). This suggests that the translation of DDX17 may be regulated by m6A in response to methionine deprivation, which could, in turn, impact the processing of its target RNA.

**Figure 5.**
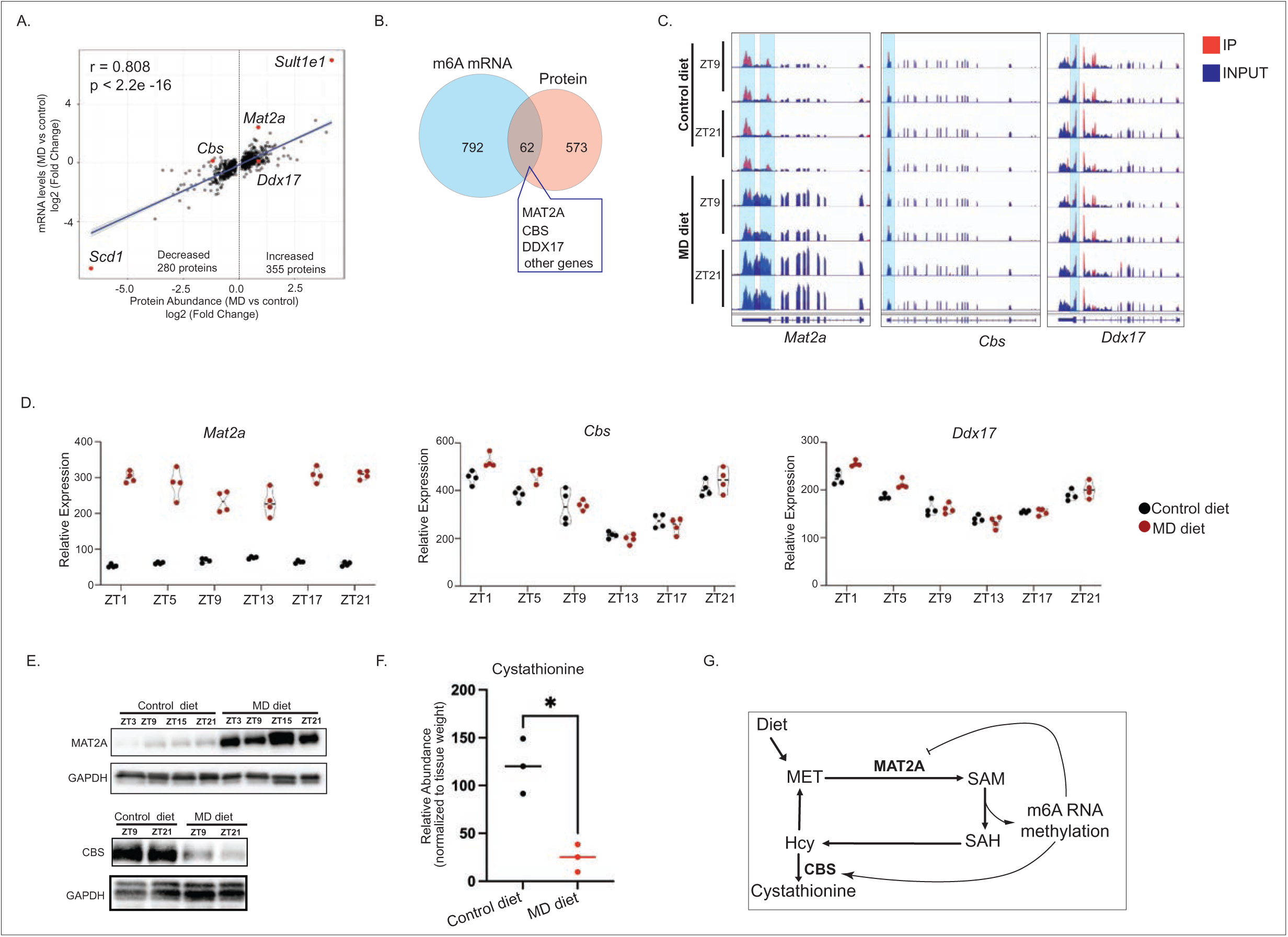
Alterations in m6A RNA modifications associated with changes in protein abundance upon MD diet. A) Scatter plot showing the correlation between mRNA expression levels and protein abundance changes upon methionine starvation. B) Venn diagram depicting the overlap between transcripts with differential m6A RNA methylation and differentially expressed proteins in the liver at ZT21. C) IGV browser tracks showing m6A RNA methylation of *Mat2a*, *Cbs*, and *Ddx17*. D) TMM-normalized gene expression values of *Mat2a*, *Cbs*, and *Ddx17*. E) Western blot depicting protein levels of MAT2A and CBS in the liver. Diet groups and Zeitgeber time are indicated above the figure. GAPDH is included as a loading control. F) The cystathionine levels in liver of mice on control diet and MD diet for three weeks (*p <0.05, two-tail t-test) G) The chart illustrates the proposed models of m6A RNA methylation in the regulation of methionine metabolism.

To further validate the results of our quantitative proteomics analysis, we measured MAT2A and CBS protein levels using immunoblotting. The western blot results confirmed that CBS exhibited lower protein levels, while MAT2A protein levels were increased in mice fed the MD diet (Figure 5E). A previous study has shown that methionine starvation leads to an increase in MAT2A levels by reducing m6A methylation and promoting RNA splicing ^18^. Consistent with this finding, we observed reduced m6A methylation levels of *Mat2a* and elevated *Mat2a* mRNA and protein levels in the liver in response to methionine starvation (Figures 5C-E).

Homocysteine, a metabolic intermediate in methionine metabolism, can either be recycled into methionine or enter the transsulfuration pathway to form cystathionine through the action of CBS. A previous study showed that a methionine-free diet causes post-transcriptional downregulation of CBS in the mouse liver ^27^. In our study, we found that the MD diet led to a reduction in CBS protein and m6A RNA modification, without affecting the mRNA level, suggesting that m6A methylation may regulate CBS translation (Figures 5C-E). Notably, in line with the reduced CBS protein levels, the cystathionine levels in liver were decreased in liver of mice fed-on MD diet (Figure 5F).

Although m6A levels of *Cbs* and *Mat2a* were both reduced by the MD diet, the changes in mRNA and protein levels between these two genes differed. This suggests that distinct m6A reader proteins recognize these transcripts and mediate different downstream regulatory outcomes. Collectively, these data demonstrate that dynamic m6A RNA methylation contributes to the alterations in mRNA and protein expression in response to methionine starvation.

## DISCUSSION

The circadian clock is tightly connected to metabolism. Disruption of the circadian clock can lead to various metabolic diseases ^28^. Conversely, metabolic interventions can reset the circadian clock ^10^. As one of the critical post-transcriptional modifications, m6A RNA modification has been shown to regulate various biological processes, including circadian rhythms and metabolic homeostasis ^29–31^. m6A RNA methylation controls circadian rhythms by regulating RNA processing, nuclear export, and translation of clock genes, including *Per2*, *Arntl*, *CK1 δ*, *Nr1d1*, and *Clock* ^12–15^. Conversely, disruption of the circadian clock through the ablation of clock genes such as *Arntl* and *Cry1/2* affects m6A modifications ^32,33^, highlighting the intricate interactions between m6A RNA methylation and circadian rhythms. In this study, we aimed to explore the role of m6A modification in the reprogramming of hepatic circadian rhythms induced by methionine starvation. Our MeRIP-seq data clearly show m6A RNA modification of several core clock genes in the mouse liver. Although the m6A levels of these clock genes are resistant to methionine starvation, we identified approximately 1,000 m6A methylation peaks exhibiting diurnal oscillations in response to methionine starvation. This suggests that m6A RNA methylation contributes to the reshaping of hepatic rhythms, likely not by targeting the core clock genes directly.

Methionine restriction leads to altered methylation by reducing SAM concentration ^10,34^. m6A RNA methylation of *Mat2a* mRNA, catalyzed by METTL16, plays a critical role in maintaining SAM homeostasis by regulating RNA splicing and stability ^18,19^. We found that, in response to methionine starvation, reduced m6A levels of Mat2a mRNA result in increased mRNA and protein expression, thereby elevating the rate of SAM synthesis. On the other hand, to compensate for methionine deficiency, reduced m6A levels in *Cbs* mRNA may impair its translation, decreasing CBS protein levels in the liver. This, in turn, could enhance the recycling of homocysteine into methionine (Figure 5G). Our findings offer new insights into the role of m6A in regulating metabolic homeostasis. However, the precise function of m6A RNA methylation in CBS translation requires further investigation.

We observed that the majority (715 out of 986) of the oscillating m6A peaks upon MD diet showed lower abundance at ZT21, which aligns with the lower serum SAM and methionine levels in mice on the MD diet at ZT21. This suggests that the de novo rhythmic m6A oscillation may be correlated with the oscillation of cellular SAM availability. SAM availability regulates the mTORC1 pathway through PRMT1 and SAMTOR ^16,17^. In the absence of methionine, SAMTOR binds to GATOR1 and inhibits mTORC1 signaling. Under methionine-sufficient conditions, SAM disrupts the SAMTOR-GATOR1 complex by directly binding to SAMTOR, which facilitates the association of PRMT1 with GATOR1, leading to mTORC1 activation ^16,17^. Notably, two recent studies reported that activation of the mTORC1 signaling pathway promotes m6A mRNA methylation ^35,36^. Therefore, the m6A oscillations observed in our study in response to the MD diet may be mediated by the mTORC1 pathway. Surprisingly, we found that *Rptor* and *Mlst8*, which encode Raptor and mLST8 in the mTORC1 complex, respectively, exhibited de novo m6A methylation oscillation in mice on the MD diet. This implies that m6A RNA methylation may also regulate mTORC1 activity in response to SAM/methionine starvation.

Although our study identifies widespread alterations in diurnal m6A RNA modification induced by methionine deprivation, several questions remain unresolved. First, our analyses establish strong associations among m6A RNA methylation, mRNA abundance, and protein expression; however, the direct causal contribution of individual m6A sites to transcriptional or translational regulation remains to be determined. In addition, future studies are needed to elucidate the mechanisms by which methionine starvation selectively induces diurnal oscillations of m6A RNA methylation in genes involved in specific signaling pathways, including the mTORC1 pathway.

## MATERIALS AND METHODS

### Animals

The animal procedures were approved by the NIEHS Animal Care and Use Committee and conducted in accordance with NIH guidelines for the care and use of laboratory animals. Six-month-old male C57BL/6J mice (JAX, 00064) were obtained from the Jackson Laboratory and housed under specific-pathogen-free (SPF) conditions on a 12-hour light:12-hour dark cycle. After a 2-week acclimatization period, the mice were fed ad libitum either an amino acid-defined control diet (Envigo, TD.01084, 0.82% methionine) or an MD diet (Envigo, TD.140119, 0% methionine) for 3 weeks. The light phase began at ZT0 (6 am, lights on) and ended at ZT12 (6 pm, lights off).

### Methylated RNA immunoprecipitation sequencing (MeRIP-Seq) and data analysis

Total RNA was extracted from snap-frozen liver with TRIzol Reagent (Invitrogen, 15596026). mRNA was purified from total RNA with PolyATtract® mRNA Isolation Systems (Promega, z5310). To fragment mRNA to a median size of ∼100 nt, the amount of 1 µl of 10X RNA Fragmentation Buffer (Life Technologies, AM8740) is added to 4 µg of mRNA in the 9 µl of nuclease-free water and incubated for 10 min at 70 °C. After adding the 1 µl of stop solution, fragmented mRNA was collected with ethanol precipitation and then resuspended the pellet in RNase-free water. A fraction of the total fragmented mRNA was reserved as input control for each sample. MeRIP was performed according to the previously reported protocol with several modifications ^37^. Briefly, 25 µl of Dynabeads™ Protein G (Thermo Fisher Scientific, 10004D) was washed with IP buffer three times (150 mM NaCl, 10 mM Tris-HCl pH 7.5, 0.1% NP-40 in nuclease-free water) and resuspended in 100 µl of IP buffer. After incubating with 3 µl of anti-m6A antibody (Abcam, ab151230) at 4 °C for 3 hours, the antibody–bead mixture was washed twice in the IP buffer. The anti-m6A conjugated beads were then incubated with 3 µg of fragmented mRNA with rotation at 4°C for 2 hours in 500 µl IP buffer with 5 µl RNase inhibitor (Thermo Fisher Scientific, N8080119). The beads were washed twice with IP buffer, twice with low-salt IP buffer (50 mM NaCl, 10 mM Tris-HCl pH 7.5, 0.1% NP-40), and twice with high-salt IP buffer (500 mM NaCl, 10 mM Tris-HCl pH 7.5, 0.1% NP-40). RNA was eluted from the beads with 250 µl of RLT buffer supplied in RNeasy Mini Kit (QIAGEN, 74106) for 5 min at room temperature. m6A-immunoprecipitated RNA was purified and concentrated using ethanol precipitation. 20 ng of untreated fragmented mRNA was used as input control. Input and immunoprecipitated RNA were reverse transcribed by SuperScript™ II Reverse Transcriptase (Invitrogen,18064014) and the sequencing libraries were constructed using the Illumina TruSeq mRNA library preparation kit. Sequencing was performed by Illumina NextSeq 500 or Illumina NovaSeq 6000 (single-end, 75 bp) at the Epigenomics and DNA Sequencing Core Facility, NIEHS.

Sequencing reads were aligned to mouse genome mm10 with STAR v2.7.9a at default parameters ^38^. m6A peaks were identified with TRESS-v.1.2.0 package and Metagene plots were generated using MetaTX-v.1.0 ^39,40^. The consensus motif of m6A peaks was detected by the HOMER motif analysis^41^. The differential m6A level was analyzed using TRESS-v.1.2.0 ^39^.

### RNA-seq data analysis

The RNA isolation and RNA-seq library preparation have been described in our previous study ^10^. The raw data we used were downloaded from GEO: GSE240881. Reads were mapped to mm10 reference genome with STAR v2.7.9a at default parameters ^38^. The differentially expressed genes were identified using EdgeR v3.30.3 at FDR < 0.01^42^.

### Mass Spectrometry-Based Proteomics with Label-Free Relative Quantification

Peptide samples were prepared using the Thermo Fisher EasyPep Mini MS sample preparation kit following the manufacturer’s recommendations. Briefly, 10 mg of liver tissue was placed in a 2 mL microtube with 1.4 mm ceramic beads (Omni International) with 200 μL lysis buffer and homogenized using a Bead Ruptor 24 (Omni International) with two 45-s pulses at 5.5 m/s with a 30 s pause between pulses. Protein content was quantified using the Bradford Assay, then proteins were reduced, carbamidomethylated, alkylated and digested with Lys-C and trypsin. After peptides were purified using spin columns, each of the samples were analyzed in triplicate using LC-MS/MS on a Q Exactive Plus mass spectrometer (Thermo Fisher Scientific) interfaced with a nanoAcquity UPLC system (Waters Corporation) equipped with a 75 µm x 200 mm HSS T3 C18 column (1.8 µm particle, Waters Corporation) and a Symmetry C18 trapping column (180 µm × 20 mm) with 5 µm particle size at a flow rate of 450 nL/min. The trapping column was positioned in-line of the analytical column and upstream of a micro-tee union which was used both as a vent for trapping and as a liquid junction. Trapping was performed using the initial solvent composition. Approximately 3 µg of peptide digest was injected onto the column. Peptides were eluted by using a linear gradient from 99% solvent A (0.1% formic acid in water (v/v)) and 1% solvent B (0.1% formic acid in acetonitrile (v/v)) to 40% solvent B over 100 minutes. For the mass spectrometry a data dependent acquisition method was employed with a dynamic exclusion time of 15 seconds and also exclusion of singly charged ions. The mass spectrometer was equipped with a NanoFlex source and a stainless-steel needle and was used in the positive ion mode. Instrument parameters were as follows: sheath gas, 0; auxiliary gas, 0; sweep gas, 0; spray voltage, 2.7 kV; capillary temperature, 275 °C; S-lens, 60; scan range (m/z) of 375 to 1500; 1.6 m/z isolation window; resolution: 70,000; automated gain control (AGC), 3 × 10e6 ions; and a maximum IT of 100 ms. For the MS/MS scans: TopN: 10; resolution: 17500; AGC 5 x10e4; maximum IT of 50 ms; and an (N)CE: 27. Mass calibration was performed before data acquisition using the Pierce LTQ Velos Positive Ion Calibration mixture (ThermoFisher Scientific). Data were processed using Proteome Discoverer also from ThermoFisher (https://www.thermofisher.com/) using a processing workflow that employed nodes for the Minora Feature Detector, Sequest HT, and Percolator. Data were searched against the mouse Uniprot sequence database using Sequest and included settings of: trypsin specificity; allowance for two missed cleavages; 20 ppm mass tolerance for MS; 0.6 Da mass tolerance for MS/MS; static modification of cysteine residues with carbamidomethylation; variable methionine oxidation; and variable modification of asparagine and glutamine deamidation. A consensus workflow containing Feature Mapper and Precursor Ion Quantifier allowed for label-free quantification.

### Immunoblotting

Frozen liver tissues were homogenized in RIPA buffer (Thermo Fisher Scientific, 89901) containing Halt Protease and Phosphatase Inhibitor Cocktail (Thermo Fisher Scientific, 1861280). Lysates were placed on ice for 30 min and then incubated at 4 °C for 90 min on a rotator. After a 15-min centrifugation at 14,000 g at 4 °C, supernatant was collected. Proteins were separated on 4-12% Bis-Tris Nupage gels (Invitrogen, NP0336BOX) and then transferred onto PVDF membrane (Bio-Rad, 162-0218) followed by incubation with primary antibodies against MAT2A (Abcam, ab186129), GAPDH (Santa Cruz, sc-32233) and CBS (Santa Cruz, sc-133154). Blots were visualized using the ChemiDoc XRS system (Bio-Rad).

### Metabolites measurement

The relative abundance of serum methionine and SAM was measured as described in our previous study ^10^. Briefly, serum proteins were precipitated and the supernatant was reconstituted in 10 mM ammonium acetate pH 5.3 for analysis after evaporation under vacuum. Methionine and SAM were separated by reverse phase liquid chromatography. Analytes were detected using tandem quadrupole mass spectrometry. Relative abundance was calculated by averaged peak area of three instrumental injections.

## Data availability

MeRIP-seq has been deposited in the NCBI database with accession number: GSE240877. Proteomics data has been deposited in the MassIVE data repository with accession number: MSV000097833.

## Author contributions

Y.L. and P.A.W. conceived and designed the study. Y.L. performed the experiments and analyzed the data. K.C. performed the animal experiments and collected tissue samples. K.L.J. and J.G.W. performed the quantitative proteomics experiments. F.B.L. and L.J.D. performed the mass spectrometry analyses for metabolite measurements. S.A.G. assisted with bioinformatics and data analysis. P.A.W. supervised the study. Y.L. and P.A.W. wrote the manuscript with input from all authors.

## Acknowledgments

This research was supported by the Intramural Research Program of the National Institutes of Health (NIH) (ES101965 to P.A.W.). The contributions of the NIH authors were made as part of their official duties as NIH federal employees, are in compliance with agency policy requirements, and are considered Works of the United States Government. However, the findings and conclusions presented in this paper are those of the authors and do not necessarily reflect the views of the NIH or the U.S. Department of Health and Human Services. We gratefully acknowledge the Epigenomics and DNA Sequencing Core Facility and Molecular Genomics Core Facility at NIEHS for assistance with NGS library sequencing.

